# An optimized live imaging and growth analysis approach for Arabidopsis Sepals

**DOI:** 10.1101/2024.01.22.576735

**Authors:** Avilash Singh Yadav, Adrienne H.K. Roeder

## Abstract

**Background:** *Arabidopsis thaliana* sepals are excellent models for analyzing growth of entire organs due to their relatively small size, which can be captured at a cellular resolution under a confocal microscope [1]. To investigate how growth of different tissue layers generates unique organ morphologies, it is necessary to live-image deep into the tissue. However, imaging deep cell layers of the sepal is practically challenging, as it is hindered by the presence of extracellular air spaces between mesophyll cells, among other factors which causes optical aberrations. Image processing is also difficult due to the low signal-to-noise ratio of the deeper tissue layers, an issue mainly associated with live imaging datasets. Addressing some of these challenges, we provide an optimized methodology for live imaging sepals and subsequent image processing. This helps us track the growth of individual cells on the outer and inner epidermal layers, which are the key drivers of sepal morphogenesis.

**Results:** For live imaging sepals across all tissue layers at early stages of development, we found that the use of a bright fluorescent membrane marker, coupled with increased laser intensity and an enhanced Z-resolution produces high-quality images suitable for downstream image processing. Our optimized parameters allowed us to image the bottommost cell layer of the sepal (inner epidermal layer) without compromising viability. We used a ‘voxel removal’ technique to visualize the inner epidermal layer in MorphoGraphX [2, 3] image processing software. Finally, we describe the process of optimizing the parameters for creating a 2.5D mesh surface for the inner epidermis. This allowed segmentation and parent tracking of individual cells through multiple time points, despite the weak signal of the inner epidermal cells.

**Conclusion:** We provide a robust pipeline for imaging and analyzing growth across inner and outer epidermal layers during early sepal development. Our approach can potentially be employed for analyzing growth of other internal cell layers of the sepals as well. For each of the steps, approaches, and parameters we used, we have provided in-depth explanations to help researchers understand the rationale and replicate our pipeline.

## Background

There is a close link between the shape of organs and the specialized function they perform. Multiple studies have shown that differential growth patterns across tissue layers generate characteristic organ shapes, which allow the organs to perform their specific functions [4–9]. For example, the twisting of plant roots increases the force generated at the tip, which aids in better soil penetration [10], while on the other hand, most plant leaves have a smooth shape that enhances photosynthesis efficiency [11]. Although the connection between organ shape and function is well known, what factors influence tissue growth leading to distinct organ morphologies is an area of active research. Therefore, to fully comprehend the organ development process, a thorough assessment of growth across multiple tissue layers is essential.

The *Arabidopsis thaliana* (henceforth Arabidopsis) sepals are the outermost green leaf-like floral organs; their smooth shape aids in properly enclosing and protecting growing flowers through development (Fig.1A, [1]). Sepals are the first organs to initiate during floral development. Like leaves, sepals have an outer (abaxial) and an inner (adaxial) epidermal layer, encasing the loosely packed mesophyll cells [1]. Being anatomically very similar to leaves, sepals serve as a great model organ for studying how plant aerial organs develop. Their small size and robustness to environmental variation allow us to study organ development at cellular resolution [1]. For live imaging, sepals are relatively easier to access compared to internal floral organs and young leaves, since removal of surrounding tissue, which is essential for imaging is less complex [1, 12]. To live image sepals from early stages, prior dissection of the older flowers off the inflorescence is imperative, since they obscure the younger flowers from the field of view (Fig. 1A, Supplementary Fig. 1). This is because flowers initiate from the inflorescence meristem continuously, following a spiral phyllotactic pattern [13–15]. This arrangement results in the younger flowers being closer and the older flowers being located further away from the meristem (Fig 1 A). Since the pedicel also elongates rapidly in an upward direction, the older flowers tend to obscure the younger flowers, and thus dissection of the older flowers becomes necessary. Once the young flowers are exposed after dissection, the sepals can be live imaged through multiple time points for growth analysis.

**Fig. 1:**
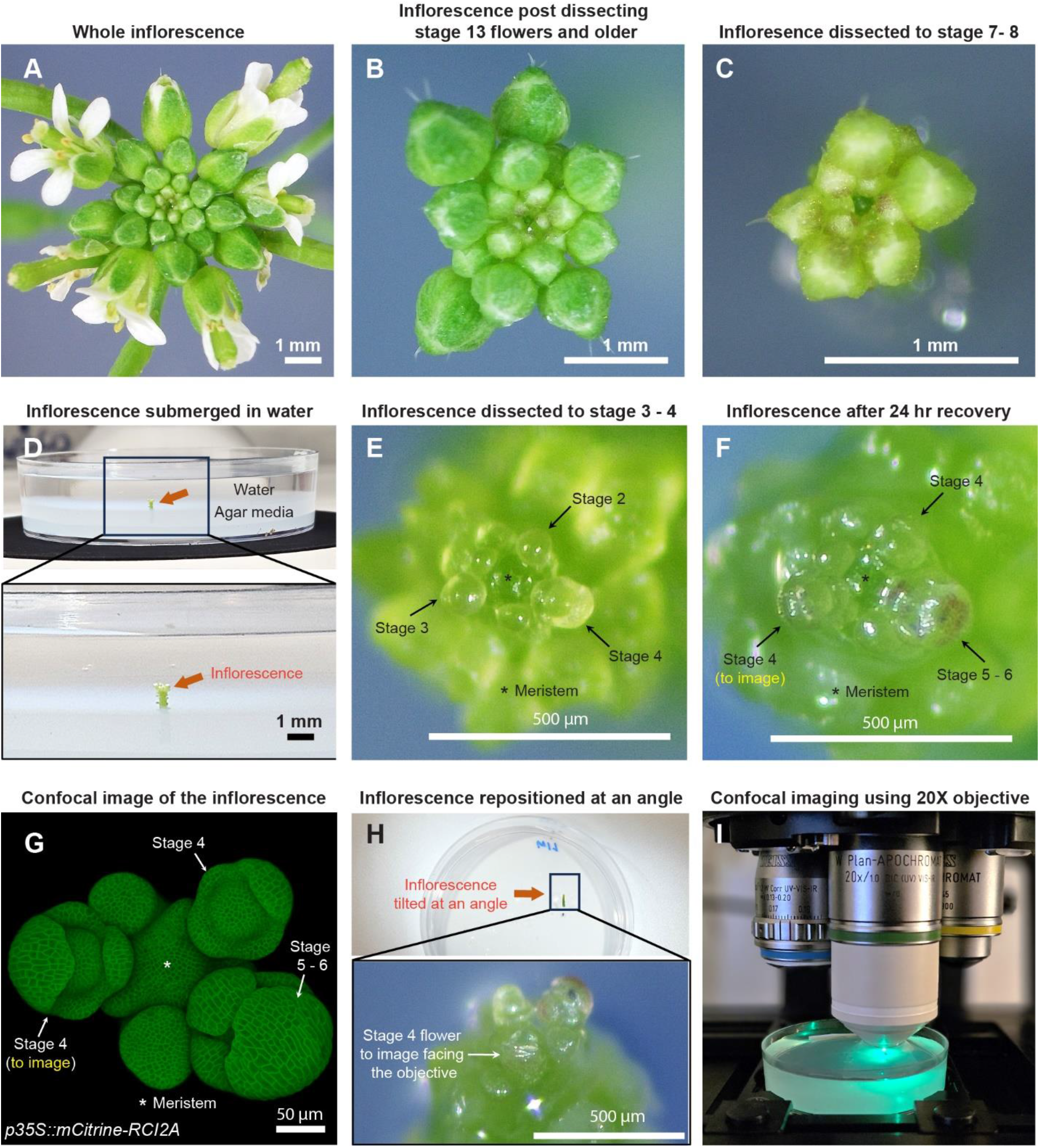
Sample preparation for live imaging sepals. **A** Whole inflorescence image of *Arabidopsis thaliana* Ler-0 ecotype harboring pLH13 (*p35S::mCitrine-RCI2A*) membrane marker. **B** Same inflorescence after dissecting flowers that are stage 13 (flower buds open with visible petals) or older. Staging performed in accordance with Smyth et. al [14]. **C** Inflorescence dissected down to flowers of stages 8 or below. Below this stage, it’s preferable to perform the dissection in water. **D** Inflorescence (from Panel C, red arrow) set in agar media containing petri plate is submerged in water for rehydration. **E** Inflorescence dissected down to flowers of stage 4 or below in water. Flowers of stages 2, 3, and 4 are indicated by arrows, with the meristem marked by an asterisk (*). **F** Inflorescence post recovery for 24 hours in growth chamber. Note the progression of flowers through the developmental stages. The stage 4 flower to be imaged is shown. **G** Confocal image of the inflorescence (panel F) under 20X magnification. **H** The inflorescence (panel F, G) is angled such the stage 4 flower faces the objective for imaging. **I** The flower being imaged under confocal microscope under 20X objective using 514 nm excitation laser.

In addition to the need for dissection, we need to live-image deeper than previous studies into the sepal tissue to track the growth of the inner epidermis. It is well established that the epidermal layers of leaves and sepals are the primary determinants of organ shape and size [16–18]. The epidermis acts as a mechanical protective layer that can constrain or promote the growth of the underlying tissues. Hence, for assessing what growth properties determine organ shape and size, we mainly focus on analyzing growth of both the outer and inner epidermal layers. Previous research on sepal morphogenesis has focused on the outer epidermis [References: Roeder 2010, Tauriello 2015, Hervieux 2016, Hong 2016, Tsugawa 2017 Development] due to technical limitations. Imaging deeper into the sepal tissue to capture the inner epidermal layer is challenging, primarily due to the following factors that hinder the penetration of light through the tissue. First, the strong autofluorescence of chlorophyll interferes with fluorophore signal detection, which affects imaging depth [19–21]. Second, extracellular air spaces in the mesophyll layer interferes with light transmission through the tissue [22, 23]. Third, the intracellular components contribute to the variation in refractive indexes which affect light penetration, thus compromising the signal-to-noise ratio [24, 25]. Optical clearing methods are routinely used to remove chlorophyll and homogenize refractive indexes of intracellular components [26–29]. However, since tissue clearing destroys its viability, such methods are inapplicable for live imaging. Overall, live imaging sepals through all tissue layers poses significant challenges, thus requiring optimization.

Alongside imaging an optimized data-driven image processing approach is required for segmentation on a curved surface of the tissue (2.5D) [2]. Segmentation in 2.5D provides a decent compromise between 2D and 3D segmentation at a reasonably low computational cost. Segmentation in 2.5D is ideal when volumetric information is not required, but the researcher intends to obtain more precise aerial measurements as compared to 2D. As 2.5D segmentation integrates the depth of the tissue, the amount of distortion typically associated with segmenting curved surfaces in 2D is largely minimized [3]. This property of 2.5D segmentation allows us to accurately capture spatial details related to the morphology of cells and tissues [30]. Also, since the epidermal layers are composed of a single layer of cells wherein the growth along the X and Y planes are comparatively much larger than in thickness (Z plane), analyzing growth in 2.5D is sufficiently informative. For highly curved surfaces like sepals, generating a 2.5D surface for the outer epidermal layers allows for accurate segmentation, lineage tracking and subsequent extraction of aerial growth properties of individual cells [31–33], since the cell depths remain relatively consistent. However, the challenge remains in generating a decent 2.5D surface for the curved inner epidermal layer which can be segmented. This challenge is posed due to the noise associated with signal from the inner epidermal layer and the variation in spatial signal intensity.

Here, we describe a detailed methodology for live-imaging Arabidopsis sepals and processing the images to track the development of individual cells on both outer and inner epidermal layers. We provide an improved and detailed procedure for floral dissection to reveal the young flowers [34, 35], with video illustrations. We describe the parameters optimized for confocal live imaging sepals across tissue layers from early developmental stages at high resolution. Using MorphoGraphX [2, 3] image processing software, we developed a technique for generating 2.5D surfaces of the inner epidermal layers to accurately segment individual cells. Finally, we discuss the parent tracking method using MorphoGraphX to follow individual cell development over time and extract growth related information. This optimized methodology serves as a comprehensive guide for researchers interested in live imaging sepal development.

## Results and discussion

### Sample preparation for confocal live imaging young flowers

For optimal dissection, we prefer six- to seven-week-old Arabidopsis plants grown in 21°C long-day conditions, which prolongs vegetative growth phase relative to 24 hours light, resulting in larger meristem sizes and thicker inflorescence axes. Dissection can either be performed while the inflorescence is on the plant or on a clipped inflorescence to allow for easier maneuvering during dissection (Supplementary Fig. 1). However, clipping the inflorescence bears the risk of faster dehydration, and thus requires quicker dissection. The choice of whether to dissect *in planta* or on a clipped inflorescence depends on the researcher.

Although the dissection techniques vary among researchers, we describe the procedure that works best for us, resulting in minimal tissue damage. For dissection, we grip the inflorescence in one hand, and use the tweezers to press the base of the pedicel and pull the flower backwards in a gentle yet swift manner (Video S1). We adopt a stage-wise dissection approach by rotating the inflorescence, starting with the oldest flowers first and gradually progressing to younger ones, all the way down to stages 7 to 8 (Fig. 1A -C, video S1, staging performed according to [14]). The inflorescence is then clipped ∼0.5 cm (about 0.2 in) from the apex, and the base of the stem is securely embedded in a live-imaging agar media petri plate (Fig. 1D). We then submerge the inflorescence in water for 3 to 4 hours to allow for rehydration, a critical step in ensuring sample viability (Fig. 1D). Following rehydration, dissection of flowers below stages 7 to 8 is performed while the inflorescence remains submerged in water. This retains sample hydration during dissection of younger flowers, as it requires increased caution and is therefore more time consuming. While using tweezers is an option, we prefer using a needle in favor of its sharper blade (see Material and Methods for specifics) and increased precision in dissection, to avoid damaging younger adjacent flowers. Dissection in water involves rotating the plate to position the flower to be dissected at a convenient angle, pulling the flower bud backwards using the needle and nipping it from the pedicel base (Video S2). These steps are repeated until we dissect down to flowers of stages 3 to 4 (Fig. 1E). Post dissection, we drain the water and transfer the sample to a fresh plate. The sample is then allowed to recover for 24 hours under initial growth conditions. Assuming no flowers were accidentally damaged during dissection, all flowers should transition through their respective developmental stages (Fig. 1E - G).

For live imaging, we choose a mid-stage 4 [14] flower (Fig. 1F, G). The inflorescence is repositioned at an angle in which the flower to be imaged directly faces the confocal objective (Fig. 1H). The sample is then submerged in 0.01% SILWET-77 (surfactant) in water solution to reduce surface tension, which helps displace the air bubbles that can obstruct imaging. Any persisting air bubbles can be removed by gently pipetting liquid solution towards air bubble. We then allow the sample to stabilize for five to ten minutes, following which the flower is imaged under a confocal microscope using a 20X water dipping objective (Fig. 1I; also see Materials and Methods for specific settings).

### Confocal live imaging to capture sepal growth

The four sepals around Arabidopsis flowers are named according to their position relative to the inflorescence meristem [33]. The ‘outer’ sepal refers to the sepal that is positioned furthest from the meristem, while the ‘inner’ sepal is the closest (usually obscured from view, Fig. 2A). The sepals on either side are referred to as the ‘lateral’ sepals (Fig.2A)[1, 33]. For live imaging, we image the outer sepal due to its accessibility (Fig. 2A). For any given sepal, the outer (abaxial) and inner (adaxial) epidermal layers are connected at the margins, enclosing the inner mesophyll layer (Fig.2B, [1]). During early sepal development, the outer and inner epidermal layers are differentially sized, with the outer epidermis exhibiting a larger surface area than the inner epidermis [9]. Anatomically, the inner and outer epidermises emerge from a single epidermal cell layer in the floral meristem and remain connected with one another, while the mesophyll consists of 2-3 loosely packed cell layers with intervening air spaces that derive from the L2 layer of the meristem (Fig. 2 C, D) [1, 36].

**Fig. 2:**
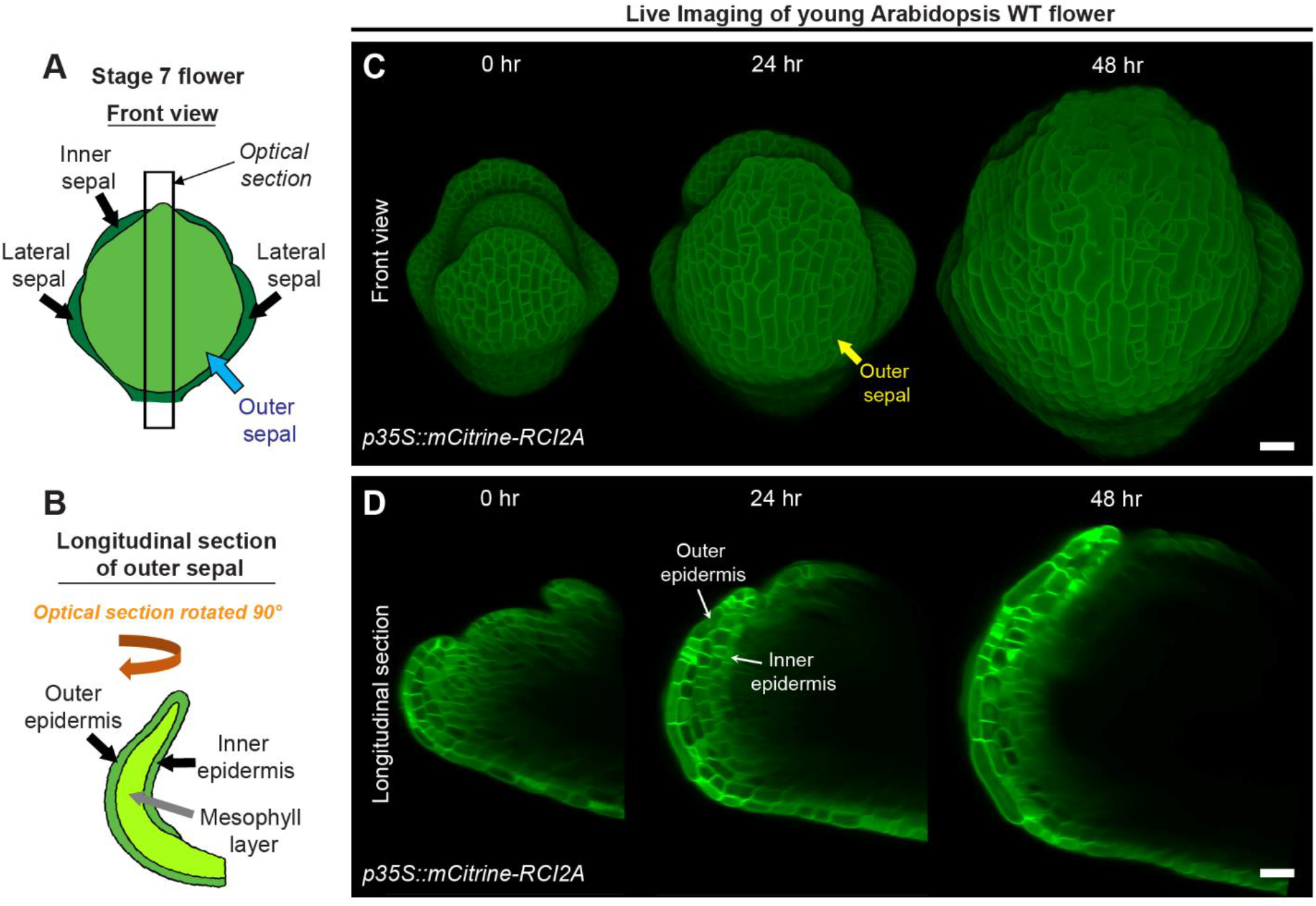
Live imaging of growing sepals. **A** Schematic diagram representing a typical stage 7 flower. The four sepals comprise an outer sepal (blue arrow) shown in light green, one inner sepal and two lateral sepals (black arrows) shown in dark green. **B** The schematic illustrates the optical longitudinal section of the outer sepal from the region marked by the black rectangle (panel A). The outer, inner epidermal layers (black arrows) in green and the mesophyll layer (grey arrow) in lime color are shown. **C** Confocal live imaging of a growing Arabidopsis flower imaged every 24 hours over a course of 48 hours with a high laser intensity for 514 nm wavelength on an argon laser at 20X magnification. The flower harbors pLH13 (*p35S::mCitrine- RCI2A*) membrane marker. The yellow arrow points to the outer sepal. Scale bar, 20 µm. **D** Longitudinal optical sections of growing flower (panel C) are shown. Note the reducing visibility of the inner layers over time. The inner and outer epidermal layers are marked (white arrows). This image series is one of the three replicates used for growth analysis in our previous study [9]. Scale bar, 20 µm.

To assess cell growth parameters underlying sepal development, we commence imaging sepals from young stages, typically from mid-stage 4 onwards (Fig. 2C). At this stage, the inner epidermis has just begun growing, and the sepals barely overlie the floral meristem [9, 14]. To visualize cell membranes, we dissect flowers from transgenic plants harboring a yellow fluorescent protein-based membrane marker (pLH13, *p35S::mCitrine-RCI2A* [37]). Capturing the membrane signal from inner epidermal cells is challenging, mainly due to the highly obscured nature of inner epidermal signal compared to the outer epidermal layer, which prompted us to optimize our imaging process (Fig. 2C, D). Firstly, we used the strong 35S viral promoter to drive membrane marker expression, which ensures robust membrane signal. Note, 35S does lead to silencing of the marker in some plants, so seedlings are screened for fluorescence before they are grown to flowering for dissection and imaging. Secondly, to enhance signal capture from deeper tissue layers without compromising sample viability, we used a high laser intensity at which the membrane signal of the outer epidermal cells was just barely saturated while that of the inner epidermal cells was unsaturated. In our Zeiss 710 confocal microscope, we achieved this with a laser intensity of 16 - 20% for the 514 nm wavelength on an argon laser. Increasing laser intensity beyond 20% on our Zeiss 710 microscope failed to improve signal intensity any further and can be detrimental to sample viability as well. Thirdly, we use isotropic voxels (x:y:z of 0.244 µm) and 1024 X 1024-pixel dimensions for obtaining high resolution images, although this considerably increases imaging time. All the three optimized parameters described above were used in combination to capture images of the quality required for image processing described in the subsequent sections. Despite these adjustments, the visibility of inner epidermal layers diminishes considerably as the flower develops (Fig. 2D). We could capture the inner epidermal layer up to stage 7, but not beyond with this approach. The factors contributing to this opacity remain undetermined, although we suspect increasing air spaces in the mesophyll tissue may either impede excitation or emission of the signal from deeper layers.

### Visualizing the inner epidermal layer of the sepal

Once the flower is imaged with the confocal microscope, the next step is reveal the inner epidermal cell layer of the outer sepal through image processing. [1] Owing to the necessity of maintaining sepal viability for live-imaging and growth tracking, the inner epidermis cannot be imaged by detaching the sepal and placing it under a coverslip. Moreover, due to the small size of the flowers below stage 8 (103-238 µm [1]), attempting to access the sepal inner epidermis by dissection is impossible without inevitably damaging the sepals. To circumvent these limitations, we employed image processing using MorphoGraphX to expose the sepal inner epidermis, by removing the voxels associated with the flower meristem regions underneath the inner epidermal layer (hereafter referred to as ‘voxel removal’). For doing so, we use the longitudinal ‘Clipping plane’ tool to visualize the longitudinal section through the flower which enables us to see the sepal internal tissues (Fig 3A-C, also see MGX user manual for more details). The depth of the Clipping plane can be adjusted using the slider under View tab/Clip 1. We recommend maintaining the Clipping plane depth at or below 20 µm, which we determined as the optimal range within which a sepal section exhibits minimum curvature. To initiate voxel removal, we position the longitudinal Clipping plane through the middle of the sepal (Fig. 3A, B). This serves as a convenient reference since the middle region of the sepal has reduced curvature compared to peripheral regions. For editing, the image is moved from the ‘Main’ Stack to the ‘Work’ Stack, using the ‘Stack/Multistack/Copy Main to Work Stack’ function. Once the Clipping plane is positioned, enabling the Clipping plane reveals the longitudinal section of the flower (Fig. 3C). Visualization of the longitudinal section can be improved by maximizing the image opacity (Main/Stack 1/Opacity) and adjusting the brightness and contrast of the view window, especially given the weak signal of the inner epidermal layer.

**Fig. 3:**
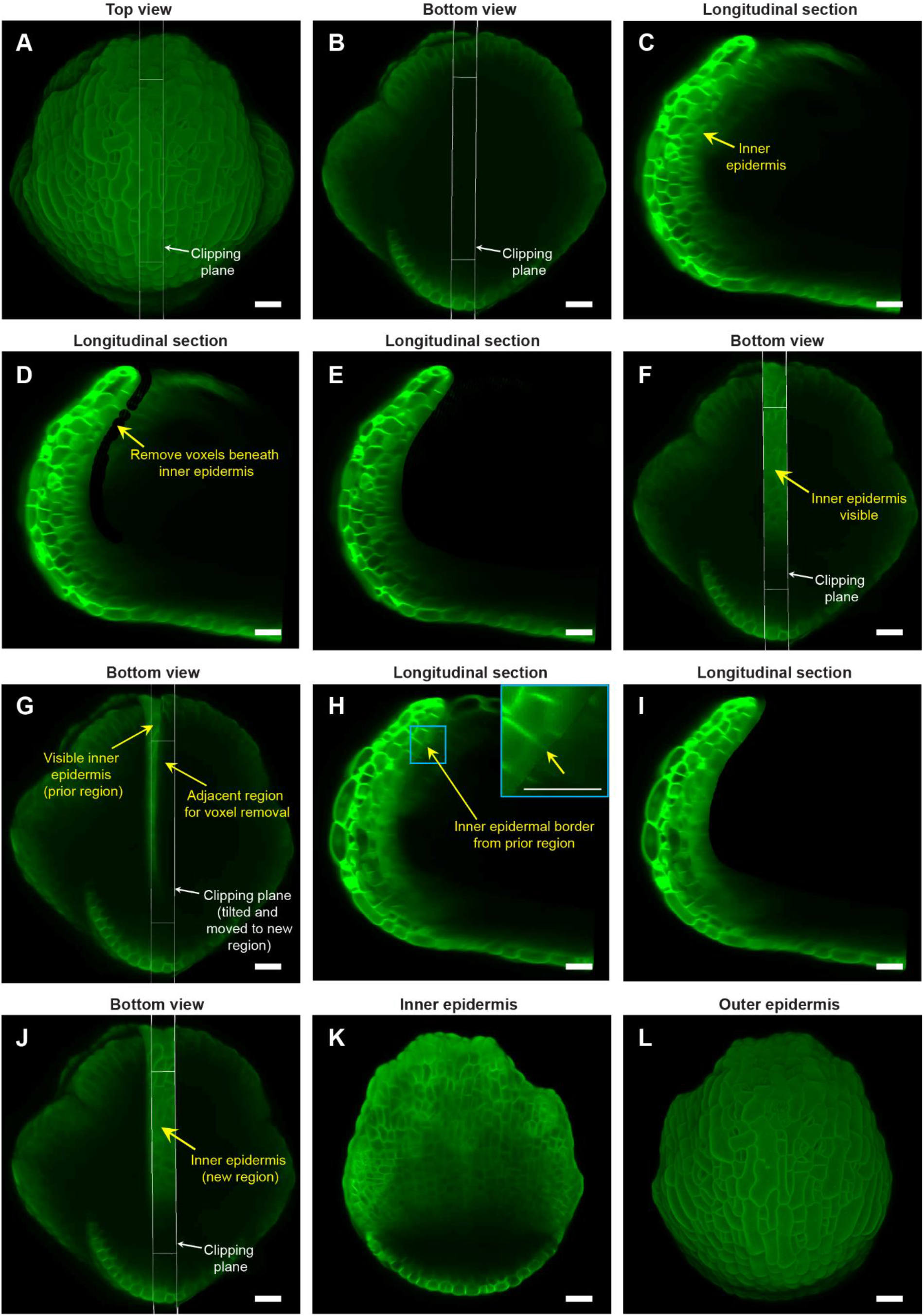
Image processing steps to expose the inner (adaxial) epidermis of the sepal. **A** Top view of the stage 7 flower imaged at the 48-hour time point of live imaging, with the longitudinal clipping plane (white rectangular box) through the center of the outer sepal blade. **B** Bottom view of the flower (panel A) with the longitudinal clipping plane positioned through the same region. **C** Longitudinal section of the flower within the clipping plane region. The inner epidermis is marked (yellow arrow). **D** Floral meristem voxels beneath the inner epidermis are removed using the voxel edit tool in MorphoGraphX [2] image processing software. **E** Longitudinal optical section post removal of all voxels beneath the inner epidermis within the clipping plane region. **F** Bottom view of the flower post voxel removal. Note the visibility of the inner epidermis compared to surrounding regions. **G** Clipping plane tilted and moved to the adjacent region on the right with some overlap with the previously edited region (panels C-E). **H** Longitudinal section displays the inner epidermal border (yellow arrow; also see inset) created by voxel removal in the previously edited region, serving as a guide for voxel removal in the adjacent region. **I** Longitudinal section post voxel removal in the adjacent region. **J** Bottom view of the flower post voxel removal of the adjacent floral meristem region. **K** Complete view of the inner epidermis of the sepal, exposed after voxel removal. **L** Outer epidermis of the sepal. Scale bars (panel A – L) represent 20 µm.

Deleting flower meristem voxels below the inner epidermis requires the use of ‘Voxel Edit’ tool (activated using the Alt-key), which allows us to remove voxels by simply left-clicking in the desired area. The radius of the ‘Voxel Edit’ tool can either be adjusted manually using the slider under View/Stack Editing/Voxel Edit Radius, or by altering the image magnification. We recommend keeping the radius of the ‘Voxel Edit’ tool below 5 µm in the initial steps for precise voxel editing. Using this small edit radius, we carefully remove voxels below the inner epidermis (Fig. 3D, video S3). Next, we zoom out to increase the Voxel Edit radius to remove voxels from the remaining internal region of that section, until the inner epidermis within the longitudinal section defined by the Clipping plane is clearly visible (Fig. 3E, F video S3). Finally, any unevenness resulting from voxel removal can be finely smoothed out to prevent irregularities in mesh surface creation in the downstream image processing steps (video S3).

To delete floral meristem voxels underneath the inner epidermis and expose it for viewing, we undertake a two-step approach. Firstly, we move the clipping plane to the adjacent section while having some overlap with the previously exposed region (Fig. 3G, video S4). Having an overlap reveals a residual border from voxel removal in the preceding section (Fig. 3H). This border serves as a guide for subsequent voxel removal in the adjacent section. This strategy of using the border as a guide ensures uniform voxel removal by eliminating the chances of under- or over- removal of voxels across discrete longitudinal sections (Fig. 3I, J, video S4). Maintaining this uniformity is critical for obtaining a smooth mesh surface and accurately projecting membrane signal in subsequent steps. Secondly, we tilt the clipping plane such that it is roughly perpendicular to the curvature of the sepal in the region from which we intend to remove voxels (Fig. 3G, video S4). ‘Voxel Edit’ deletes voxels all the way through the thickness of the section. Since the sepal is a curved tissue, the clipping plane must be perpendicular so that voxels associated with the inner epidermal layer on the opposite side are not accidentally deleted.

The two steps (tilting and moving the clipping plane for voxel removal) are repeatedly performed throughout the flower in a longitudinal section-by-section manner, until we achieve a complete view of the inner epidermis (Fig. 3K, video S4). Note that all the steps described above are error-prone; whether voxel removal was optimally performed or not can only be assessed later, based on how good the membrane signal projection is on the mesh surface (described in the next section). Therefore, saving our progress regularly is a smart practice that allows us to revert to any previous versions if any anomalies arise, without having to start all over. To save the current image version, we simply copy the image in the ‘Work’ stack back to the ‘Main’ stack using the process ‘Stack/Multistack/Copy Work to Main Stack’ and then save the stack. Overall, the steps described above allow one to view both the inner and outer epidermis of growing sepals (Fig. 3K, L) without the need for physically dissecting the sepal. Although we illustrate the ‘voxel removal’ steps using the 48-hour time point, the same can be applied to visualize the inner epidermis for the other time-points. While we use it for obtaining the sepal inner epidermis, the technique described above can also be applied for exposing deep cell layers of other floral organs such as petals or stamens, provided the sepals are removed to facilitate imaging of internal floral organs.

### Image processing steps for cell segmentation

After the sepal inner epidermis is exposed, we then proceed with cell segmentation, which is a computational technique for demarcating individual cells that can be used for extracting cellular features [38]. We segment the cells on a computational 2.5D mesh surface that represents the sepal morphology. The mesh comprises triangles and segmenting the mesh into discrete cellular units involves labeling the triangle vertices [3]. The methodology for generating a 2.5D mesh surface is outlined in the MorphoGraphX user manual. Here, we describe how to optimize the parameters for generating a 2.5D mesh surface specifically for the epidermal layers of the sepal. Note that, while we detail the parameter optimization specific to this study, the image processing steps need to be optimized for each individual experiment depending on the type and quality of the images.

To generate the inner epidermal mesh surface, the initial image orientation which places the inner epidermis at the bottom of the stack needs to be inverted along the Z-direction (Fig. 3K, 4A). Reorienting the image stack by running ‘Stack/Canvas/Reverse Axes’ positions the inner epidermis at the top, which informs MorphoGraphX to create the mesh surface for the inner epidermis. To attenuate image noise which could interfere with inner epidermal shape extraction in the subsequent steps, we run the process ‘Stack/Filters/Gaussian Blur Stack’ with X/Y/Z sigma (µm) values ranging between 2 - 2.5 (Fig. 4B). While lower sigma values (<=1) are generally preferred, images with weaker membrane marker signal requires more blurring, which minimizes the occurrence of holes when we extract the global shape. Once the image is blurred, we can extract the global shape of the inner epidermis by running ‘Stack/Morphology/Edge Detect’, with a threshold value ranging between 2000-3000 (Fig. 4C). While a higher threshold value typically yields a shape closely resembling the image, this usually requires strong cell membrane signals. Weak signal strength (as is the case for the inner epidermis) leads to holes in the structure if higher threshold values are used. Although small holes can be corrected by running ‘Stack/Morphology/Closing’, attempts to fill large holes result in structural distortions.

**Fig. 4:**
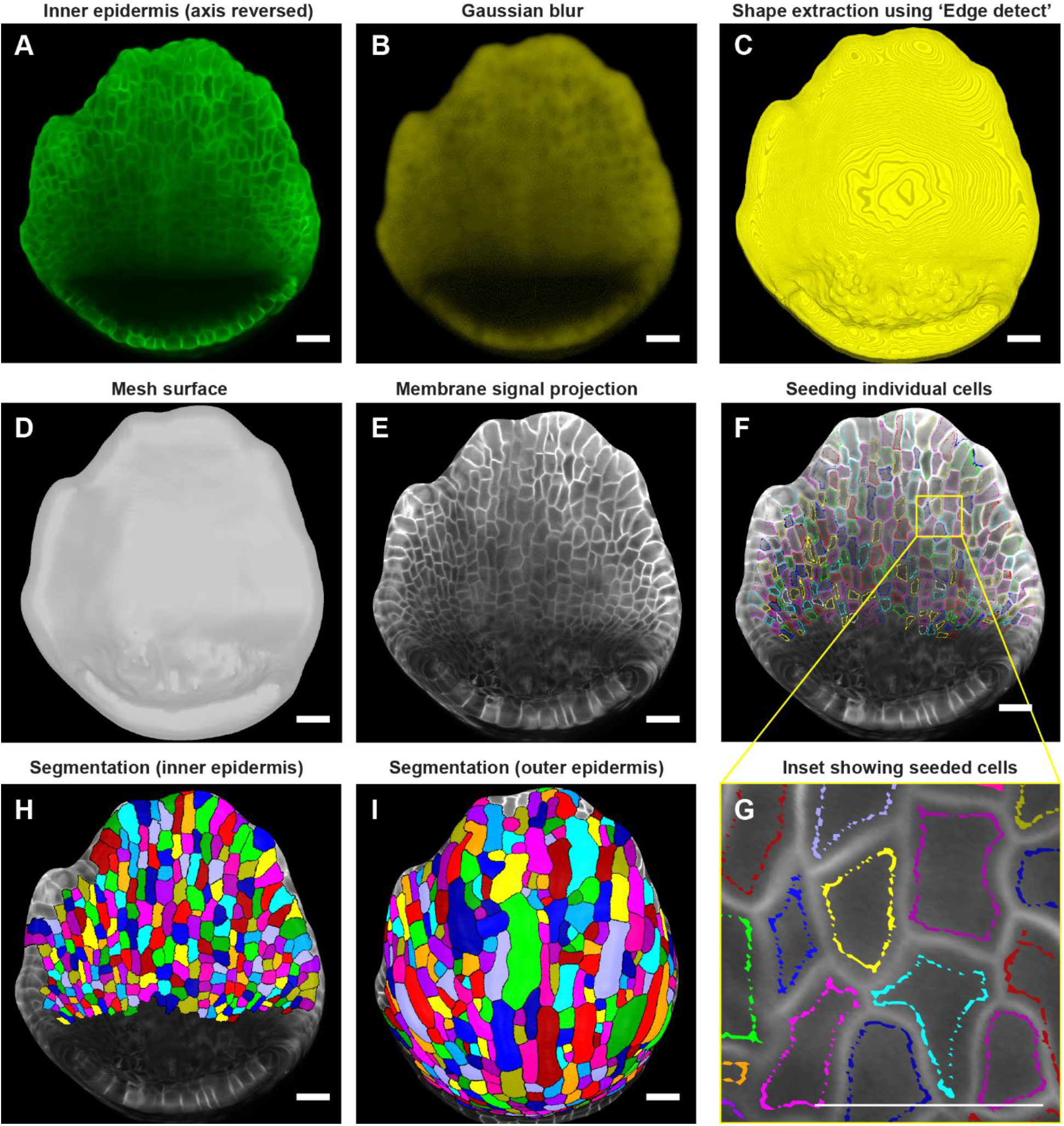
Downstream image processing for cell segmentation. **A** MorphoGraphX [2] processed image of the inner epidermis after reversing the axis along the Z direction. **B** Inner epidermal image stack post running Gaussian Blur filter. **C** Extraction of the global shape of the inner epidermis using MorphoGraphX [2] process “Edge detect”. **D** Mesh surface generated using “Marching Cubes Surface” followed by smoothing and subdividing the mesh. **E** Plasma membrane marker signal projected onto the mesh. **F** Cells seeded (labeled) throughout the inner epidermis. Colors represent individual labels. **G** Inset (panel F) demonstrates how the cell label follows the cell shape. Note that the labeling close to the cell boundary is essential for accurate segmentation especially when the membrane signal is weak. Scale bar, 2 µm. **H-I** Watershed segmentation of **H** Inner epidermal cells and **I** Outer epidermal cells. Scale bars across panels A- I represent 20 µm.

Following global shape extraction, we generate a 2.5D mesh surface by running ‘Mesh/Creation/Marching Cube Surface’ with a cube size of 3 µm (Fig. 4D). Irregularities in the mesh are then smoothed by running ‘Mesh/Structure/Smooth Mesh’ with 10 passes. We usually avoid smoothing the mesh beyond 10 passes as it can shrink the mesh, which in turn can distort membrane signal projection. To enhance mesh resolution, we increase the number of vertices by running ‘Mesh/Structure/Subdivide’. In this case, optimum resolution is typically achieved with two iterations of subdividing, beyond which additional subdividing increase the file size of the mesh (and in turn, computational time for processing) without any discernable improvement in membrane signal resolution. Next, we project the membrane signal onto the mesh using ‘Mesh/Signal/Project Signal’ (Fig. 4E). Projecting the signal with minimum and maximum projection distance of 3 and 8 µm respectively below the mesh is usually sufficient, although minor adjustments may be necessary. Typically, the lower the ‘Edge Detect’ threshold values used to extract the global shape, the deeper in microns the image needs to be projected on the mesh surface to capture the membrane signal. We then manually seed the cells using the ‘Mesh tools bar/Add new seed’ tool, activated using the Alt-key (Fig. 4F). To prevent labels bleeding into neighboring cells during segmentation, especially given that the inner epidermal cell membrane signals are weak, care is taken to seed close to the cell membrane outlining the cell shape (Fig. 4G). We then perform cell segmentation by running the process ‘Mesh/Segmentation/Watershed Segmentation’, which propagates the respective labels of individual cells (Fig. 4H). In the final step, we verify whether the segmentation of each cell accurately represents the cell morphology. If any discrepancies are found, we delete the segmentation of the given cell and its adjacent cells by filling it with an empty label, using ‘Fill Label’ from ‘Stack tools’, and then re-seeded and re-segment the cells. Once we are satisfied with the segmentation, we run the process ‘Mesh/Cell Mesh/Fix Corner Triangles’ which ensures no vertices in the mesh are unlabeled, as it interferes with the parent tracking process. Note, MorphoGraphX also allows for automatic seeding and segmentation [3]. This approach is useful for segmenting individual cells in larger samples, such as leaves or fully mature sepals, particularly when the cell boundaries are well defined. However, this approach also results in under or over segmented cells which need to be manually corrected. In this case, where the number of cells is relatively few, and the membrane signal of the inner epidermal cells is weak, manual seeding is more efficient and accurate.

Due to a considerable disparity in signal-to-noise ratio, the surfaces for the inner and outer epidermis need to be created separately to achieve optimal membrane signal projection. To segment the outer epidermal cells from the same sepal, we use the sepal image prior to reorienting the stack. While the same processes described above for creating a surface for the inner epidermis can be followed, the parameters need to be modified to account for the enhanced membrane signal intensity of the outer epidermal cells. Briefly, we run ‘Gaussian blur’ to blur the stack with X/Y/Z sigma (µm) values of 1-1.5, followed by ‘Edge detect’ for global shape extraction with threshold values of 9,000-10,000. The parameters for creating the mesh surface, smoothing, and subdividing the mesh remain consistent with those described for the inner epidermis. Optimal signal projection is typically obtained with min. and max. distances of 2 and 4 µm, respectively. Cells are then carefully seeded according to cell shape, followed by segmentation using the watershed algorithm (Fig. 4I). Taken together, the steps described are employed to segment cells on both the inner and outer epidermal layers across time points.

### Parent tracking cells through multiple time points

To determine the cell lineages over time, we use the ‘Parent tracking’ process (refer to MorphoGraphX user manual for more details), which involves linking the daughter cell labels (in the later time point (T2)) with their respective parent cell labels (in the earlier time point (T1). This process allows us to track how cells grow and divide over time (Fig. 5 A, B). For parent tracking, the segmented T1 mesh and T2 meshes are loaded into ‘Stack 1’ and ‘Stack 2’ respectively (video S5). For the T1 mesh, we deselect ‘Surface’ in the ‘Stack 1’ view window, check ‘Mesh’ and choose ‘cells’ from the ‘Mesh’ dropdown menu to retain only the segmented cell outlines. For the T2 mesh, we activate ‘Parents’ under ‘Stack 2’ view window. We prefer distinct colors for segmented cell outlines in the T1 and T2 meshes to avoid confusion, options for which are available under ‘Miscellaneous tools bar/Color Palette’.

**Fig. 5:**
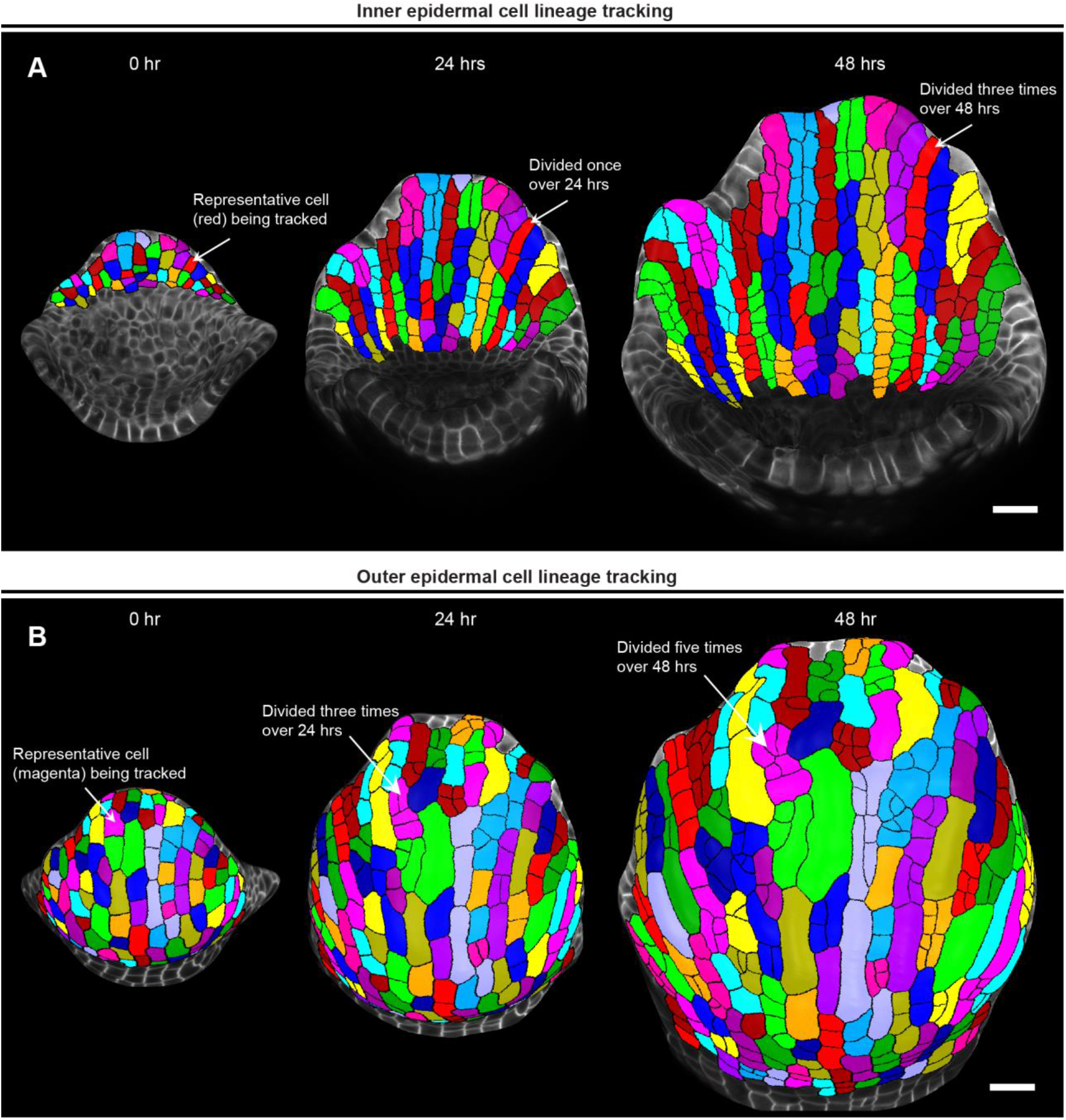
Parent tracking to quantify cell growth properties. **A** MorphoGraphX [2] generated meshes of growing inner epidermis showing segmented cells parent tracked over 48 hours. The arrow shows a representative cell (red) that divided once to generate two daughter cells in 24 hours, and thrice to generate four daughter cells in 48 hours. Scale bar, 20 µm. B Meshes showing segmented cells of the outer epidermis parent tracked over 48 hours. The arrow points to a representative cell (magenta) that has divided thrice times over 24 hours and an additional two times over 48 hours. Note the development of meristemoid cell in this lineage, which will eventually differentiate into a stoma. Scale bar, 20 µm.

To initiate the ‘Automatic parent labeling’ process in MorphoGraphX, manual parent labeling of a few daughter cells, ideally distributed uniformly across the T2 mesh, is necessary. These manually parent labeled cells act as landmarks for auto-parent labeling the remaining daughter cells (Supplementary fig S2, video S5 and S6). To transfer parent labels, we superimpose the T1 mesh onto the T2 mesh surface such that the parent cell (in T1 mesh) selected for label transfer overlies its corresponding daughter cells (in T2 mesh) (video S5). Using the ‘Mesh tools/Grab label from other surface’ tool, we then simply click within the parent cell region to transfer the label. For manual parent labeling, we prioritize labeling those daughter cells in T2 mesh that are easily recognizable based on their morphological similarity to the parent cells in T1 mesh (video S5). Conceptually, the cell neighbors do not change despite the cells undergoing growth and division. Based on this concept, the daughter cells which can be easily recognized based on their morphological similarity to the parent cell can be used as references to parent label additional daughter cells at distant locations throughout the T2 mesh. We then run the process ‘Mesh/Deformation/Morphing/Set Correct Parents’ to set these manually parent labeled cells as landmarks for auto parent labeling, which is then performed by running ‘Mesh/Deformation/Mesh 2D/Auto Parent Labelling 2D’.

While ‘Auto Parent Labelling 2D’ labels the majority of the daughter cells, the cells that are left out need to be manually parent labeled (video S6). Finally, we verify whether all the parent labels have been assigned correctly by running the ‘Mesh/Cell Axis/PDG/Check Correspondence’ process on the T1 mesh (video S6). This generates a heatmap for both time points, allowing us to visually assess whether cell junctions correspond correctly between T1 and T2 meshes (video S6). The cells with accurately matched junctions are highlighted in blue, while those with errors are highlighted in red. These discrepancies typically arise from either segmentation errors or incorrect assignment of parent labels which need to be rectified manually, with the objective of having all the cells pass the correspondence check. Following this step, keeping Stack 2 active, we save the parent labels as a .csv file by running ‘Mesh/Lineage Tracking/Save Parents’. Usually, we parent track the cells across two consecutive 24-hour time points and save the parent labels for each pair of consecutive time points. In this case, we parent tracked cells from 0-to-24-hour and 24-to-48-hour time point for both outer and inner epidermis and save the parent labels as .csv files. For parent tracking through multiple time points (Fig. 5), we use the .csv files to run a custom python script (combinelineages.r [39]) to correlate the parent labels, followed by manual correction if needed. This allows us to trace individual cells from the initial time point (0 hour) up to the final time point (48 hour) (Fig. 5).

Parent tracing cells on the outer and inner epidermal cells reveals an interesting pattern of cell growth and division. We found that almost all the inner epidermal cells divide vertically along the longitudinal axis of the sepal (Fig. 5A). While cells on the outer epidermis also exhibit cell division along the longitudinal axis, several cells divide along the transverse direction (along the width) as well (Fig. 5B). We also see multiple instances of asymmetric cell divisions, wherein the smaller sister cells can be predicted to differentiate into stomata. Additionally, since giant cell specification occurs early during development [18], cells that cease division eventually differentiate into giant cells. Overall, once we have successfully parent tracked the cells, we can extract a multitude of growth related information for each cell lineage, which includes growth rate, direction of growth, cell division rate etc. to name a few, as previously shown [9].

### Conclusions

Our optimized methodology offers a seamless workflow from sample preparation to imaging and image processing for analyzing early sepal development. Despite the faint nature of membrane marker signal from the inner epidermal layer, our pipeline enables reliable parent tracking of cell lineages. This enables us to bypass the need for sepal dissection, thus preserving sample integrity. For sample preparation, our improved protocol employs a simple, two-step approach; initial dissection of flowers off the inflorescence using tweezers, followed by a more precise dissection of remaining flowers using a needle while the sample is submerged in water. We detail the requirements for live imaging sepals across tissue layers under a confocal microscope. We explain how to optimize the imaging parameters to allow for high-quality images to be obtained without affecting sample viability and growth. For visualizing the inner epidermis of the developing sepals, we provide a detailed description of the computationally inexpensive ‘voxel removal’ technique. This technique involves the simple, yet careful use of the ‘Voxel Edit’ and ‘Clipping Plane’ features in MorphoGraphX. Given the differences in membrane signal strength between epidermal layers, we provide justifications for the parameters used in each image processing step to generate 2.5D meshes for segmenting inner as well as outer epidermal cells. Finally, we describe the parent tracking process across time points with video illustrations. We believe our methodology will be of valuable use for researchers studying cell growth dynamics of inner plant tissue layers in 2.5D, particularly in cases where the signal strength of inner tissue layers is weak.

## Materials and methods

### Plant material and growth conditions

In this study, we used the *Arabidopsis thaliana* Landsberg *erecta* (L*er*-0) ecotype plants, expressing the pLH13 (*p35S::mCitrine-RCI2A*) cell membrane marker. For imaging and subsequent analysis, plants of the T2 generation were used. The seeds were initially planted on 1/2 MS media plates (comprising 1/2 MS salt mixture, 0.05% MES w/v, 1% sucrose w/v, pH ∼ 5.8 adjusted with KOH, and 0.8% phytoagar). The media was supplemented with Hygromycin (GoldBio) at a concentration of 25 µg/ml to select transgenic containing the pLH13 construct as previously described [9]. For selection in plates, the seeds were sterilized by washing the seeds twice with a sterilization solution containing 70% ethanol + 0.1% Triton X-100, followed by two additional washes with 100% ethanol. Sterilized seeds were then dried on filter paper (Whatman 1), spread evenly on the ½ MS media hygromycin plates, and stratified at 4°C for three days. Following stratification, the plates were transferred to a growth chamber (Percival-Scientific), maintained under long-day light conditions (16 hours fluorescent light/8 hours dark) at a light intensity of approximately 100 µmol m^-1^s^-1^ and a relative humidity of 65%. The seedlings were grown on plates for 1.5 to 2 weeks until the transgenic plants that survived Hygromycin selection were clearly distinguishable from the non-transgenic counterparts. The transgenics were subsequently transplanted into pots (6.5×8.5 inches) containing soil mixed with Vermaculite (LM-111), with one plant per pot to facilitate healthy vegetative growth. For live imaging, the samples were grown in ½ MS media containing 1% sucrose and 1.2% w/v phytoagar, supplemented with 1X Gamborg Vitamin mix (containing final concentration of 100 µg/ml myo-Inositol, 1 ng/ml Nicotinic Acid, 1 ng/ml Pyridoxine HCl, and 1 ng/ml Thiamine HCl [40]) and 1000-fold diluted plant preservative mixture (PPM™, Plant Cell Technology).

### Schematic diagrams

The schematic diagrams as shown in Fig. 2 A, B were drawn by hand using the Notability application in Apple iPad (model A1701).

### Dissection

Primary inflorescences were clipped off six- to seven-week-old plants for dissection. For dissecting the inflorescences down to flower stages 7 to 8, Dumont tweezers (Style 5, Electron Microscopy Sciences) were used. For subsequent dissection of the flowers down to stages 3 to 4 in water, we used BD Micro-FineTM IV needle (BD U-100 Insulin Syringes, 0.35mm (28 - gauge needle) and 12.7 mm [1/2 in.] length)). Dissections were performed using either Zeiss Stemi 2000 or Stemi 508 stereo microscope.

### Capturing inflorescences and whole plants

Photos of the whole plant, clipped inflorescence, and the dissected inflorescence embedded in plates containing live imaging media were taken with Samsung Galaxy S23+ phone (1848 x 4000, 7 MP). Microscopic images and videos of the inflorescence dissection were captured with an Accu-Scope Excelis 4K UHD camera mounted on Zeiss Stemi 508 stereo microscope.

### Confocal imaging

The Zeiss LSM 710 confocal microscope, equipped with inverted AXIO Examiner.Z1 was used for live-imaging the sepals using a 20X water-dipping objective (Plan-Apochromat 20x/1.0 DIC D=0.17 M27 75mm). The microscope is controlled by the Zeiss ZEN user interface software (ZEN 2012 SP5 FP3 (64 bit) version 14.0.27.201) which is installed in a 64-bit Windows 10 system (CPU - Intel Xeon(R) Gold 5222 @3.80 Hz, RAM - 64.0 GB). For the 0- hour timepoint, the sepal of stage 4 flower was imaged with the following settings: x: y: z spatial scaling of 0.244 µm, image dimensions of 1024 x 1024 x 743 pixels (8-bit), a zoom factor of 1.7, averaging = 1, excited with argon laser at 514 nm with 16% intensity and a master gain of 485. For the 24-hour timepoint, the imaging settings were the same, except for image dimensions, which were 1024 x 1024 x 885 pixels (8-bit), and a master gain of 505. For the 48-hour time point, the sepal was imaged with image dimensions of 1024 x 1024 x 895 pixels (8-bit) and a master gain of 513. Across all time points, the emission for mCitrine was collected between the range of 519-596 nm. The images were saved in .lsm file formats.

### Image processing

Image processing in this study was performed using MorphographX 2.0 r1-294 image processing software [2, 3], installed in an Ubuntu 20.04.4 LTS (Focal Fossa) 64-bit operating system. The system has NVIDIA Cuda driver version 11.40 installed and is equipped with the following hardware specifications: CPU - Intel (R) CoreTM i7-5930K @ 3.50GHz with 12 cores, RAM - 64 GB, and GPU - NVIDIA GeForce GTX TITAN X with a capacity of 12 GB. Our system specifications met the requirements recommended for the MorphoGraphX software, which are 32-64 GB RAM and an NVIDIA graphics card.

For viewing and processing the images in MorphoGraphX, the raw images in .lsm file format were converted to .tiff format using Image J distribution Fiji [41]. In general, we import the first image in ‘Stack 1’ view window under ‘Main’, and any subsequent images in ‘Stack 2’. For taking screenshots of the flowers, the extraneous parts of the image not associated with the flower (such as dissected parts of the inflorescence and/or other flowers) were removed using the ‘Voxel Edit’ tool. All screenshots were captured using the ‘Save screenshot’ tool and saved as .png image files. The videos were captured using the ‘Record Movie tool’. Briefly, prior to video capturing, the images/meshes were pre-positioned in the view window and then rotated or moved in the view window as required, with the ‘Record Movie’ tool activated. The ‘Record Movie’ tool captures these movements by generating a series of two-dimensional .tiff files. To produce the videos, the .tiff file series were imported into Fiji using ‘File/Import/Image Sequence’ and were subsequently saved as uncompressed .avi video files. Further video processing, including lossless compression, labeling, and annotation, was performed using Adobe Premiere Pro 2023.

## Availability of data and materials

The images and the meshes for all the three replicates for the WT are publicly available at Open Science Framework under CC-By Attribution 4.0 International license. The images of the WT flowers, as well as the final edited images can be accessed at https://doi.org/10.17605/OSF.IO/UMW9B. The images shown in this manuscript correspond to WT replicate 2. The meshes corresponding to WT replicate 2 can be accessed under Figure 3.zip file at https://doi.org/10.17605/OSF.IO/P5Q39.

## Supporting information

Video S1: Dissecting Arabidopsis flowers off the inflorescence using tweezers.

Video S2: Dissection of Arabidopsis flowers in water using a needle.

Video S3: Removing voxels beneath the inner epidermis.

Video S4: Sequential voxel removal beneath the sepal inner epidermis in longitudinal sections using the clipping plane to expose the inner epidermis.

Video S5: Manually assigning parent labels to inner epidermal cells.

Video S6: Semi-automatic parent labeling.

## Acknowledgements

We thank Isabella Rose Burda, Shuyao Kong and Michelle Heeney for their comments on the manuscript. We thank Richard Smith (John Innes Centre, UK) for his advice on image processing.

## Funding

Research reported in this publication was supported by the National Institute of General Medical Sciences of the National Institutes of Health under Award Number R01GM134037 to AHKR. ASY is also supported by the 2023 Sam and Nancy Fleming Research Fellowship. The content of this study is solely the responsibility of the authors and does not necessarily represent the official views of the National Institutes of Health and other funders.

## Contributions

ASY and AHKR conceptualized the study. ASY generated transgenics, designed the experiments, collected, and processed the images. ASY and AHKR wrote the manuscript. All authors read and approved the final manuscript.

## Supplementary Figures

**Supplementary Fig. 1:**
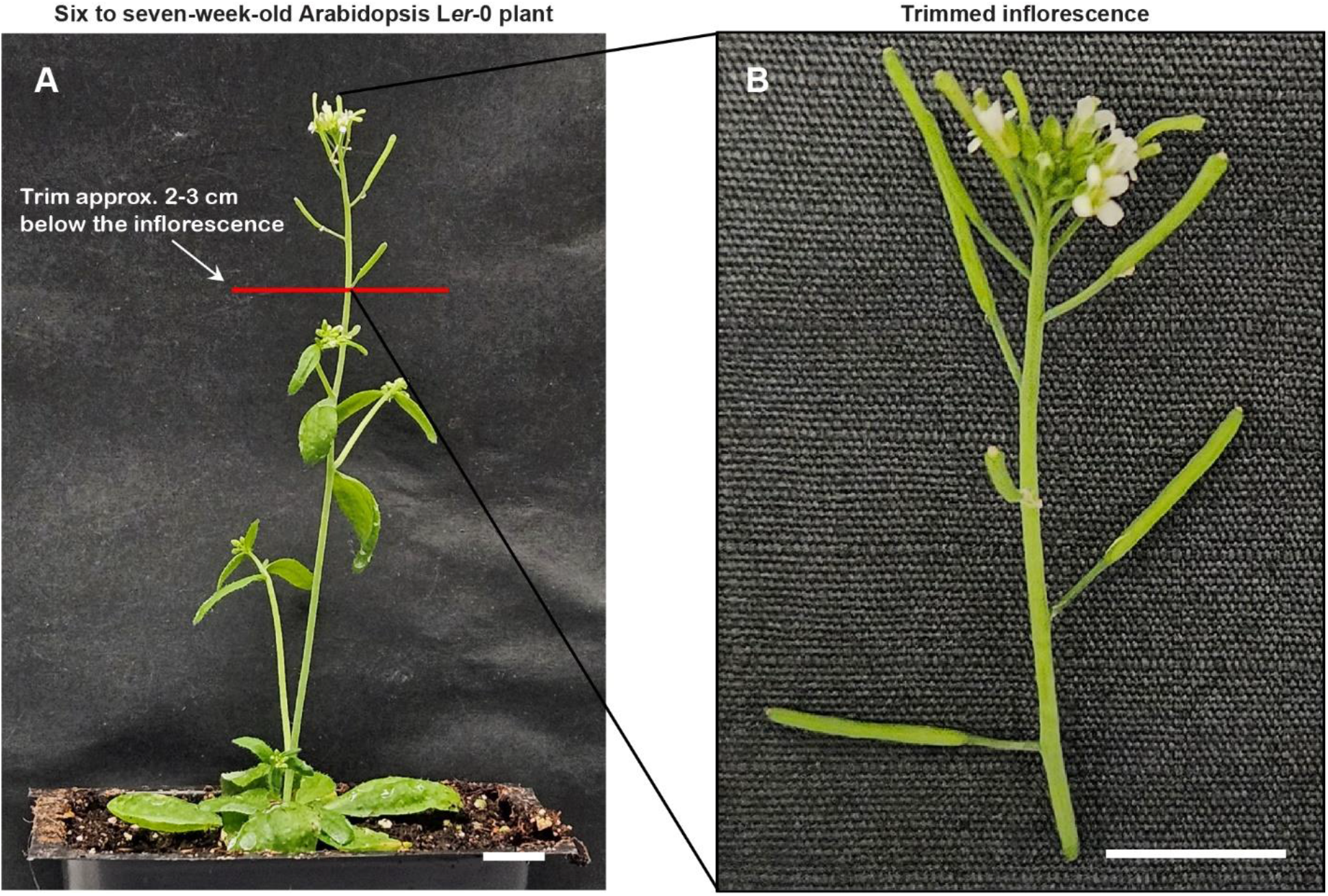
Trimmed inflorescence for dissection, associated with main figure 1. **A** Whole plant image of six to seven-week-old Arabidopsis L*er*-0 ecotype grown in long-day conditions (16-hr light/8-hr dark). **B** Inflorescence post-trimming, which has been cut approximately 2-3 cm below the inflorescence apex (red line) for dissection. Scale bar, 1 cm.

**Supplementary Fig. 2:**
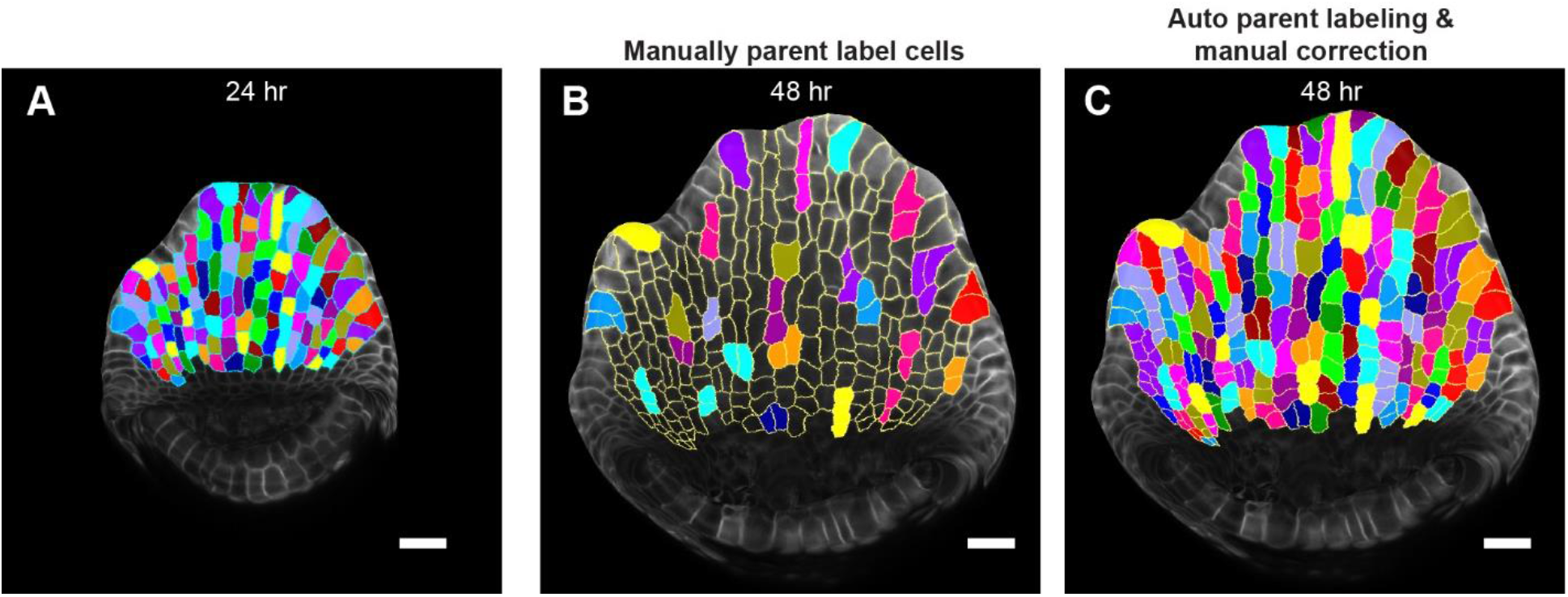
Semi-automatic parent tracking, associated with Main figure 5. **A** MorphoGraphX generated mesh for the inner epidermis at 24-hour time point showing segmented cells. Colors represent individual cells. **B** Inner epidermis mesh at 48-hour time point, wherein segmented cells are being parent tracked to cells in 24-hour time point (panel A). Colors correspond to parent cells in the 24-hour time point mesh. Note that manually parent labelled cells are distributed across the mesh, which greatly aids automating parent labelling. **C** Cells in 48-hour time point parent labelled using automatic parent labeling, followed by manual correction. Scale bar, 20 µm.

## Supplementary video legends

**Video S1:** Dissecting Arabidopsis flowers off the inflorescence using tweezers. The flowers are dissected by pressing the pedicel using tweezers while having the inflorescence securely held in one hand. As the flowers are being dissected, the inflorescence is rotated gradually such that the older flowers are dissected first, followed by younger flowers.

**Video S2:** Dissection of Arabidopsis flowers in water using a needle. The inflorescence, comprising flowers of stages 8 and below, is anchored in agar media while being submerged in water to prevent dehydration during dissection. The plate is rotated such that the flower to be dissected is at a convenient angle. Older flowers are dissected first, followed by the younger flowers.

**Video S3:** Removing voxels beneath the inner epidermis. Enabling the clipping plane (positioned longitudinally) reveals the longitudinal section of the flower. The ‘Voxel Edit’ tool is used to carefully remove voxels beneath the inner epidermis, without removing voxels associated with the inner epidermis.

**Video S4:** Sequential voxel removal beneath the sepal inner epidermis in longitudinal sections using the clipping plane to fully expose the inner epidermis. The video shows the 3D dimensional view of a typical Arabidopsis stage 7 flower, with cells on both epidermal layers of the outer sepal segmented. To expose the inner epidermis, voxels within a given longitudinal section defined by the clipping plane are removed. The steps for voxel removal involve tilting and moving the clipping plane to the adjacent regions, resulting in a complete view of the inner epidermis. The 3D image of the inner epidermis can then be used to generate a mesh surface onto which the membrane signal is projected for segmenting the cells.

**Video S5:** Manually assigning parent labels to inner epidermal cells. In this video, the inner epidermal meshes from earlier (T1, 24-hour) and later (T2, 48-hour) time points are loaded onto Stack 1 and Stack 2, respectively. To assign parent labels, the T1 mesh surface is unchecked, leaving only segmented cell outlines visible. With parents active for the T2 mesh, the T1 mesh is overlayed onto the T2 mesh, such that the parent cells lie above their corresponding daughter cells. Parent labels are assigned by clicking within the parent cell area.

**Video S6:** Semi-automatic parent labeling. In this video, 24-hour timepoint (T1) mesh and 24-hour timepoint (T2) mesh are loaded onto Stack 1 and 2 respectively. A subset of cells in the T2 mesh are manually parent labeled, which serves as references for MorphoGraphX process ‘Auto Parent Labeling 2D’ to label the remaining cells. Cells that were not auto-parent labeled were labeled manually. The accuracy of parent labeling was checked by running the process ‘Check Correspondence’ which checks whether cell junctions between T1 and T2 mesh correspond correctly.

## Notes

https://doi.org/10.17605/OSF.IO/UMW9B

https://doi.org/10.17605/OSF.IO/P5Q39

### Competing Interest Statement

The authors have declared no competing interest.

